# Multi-strategies embedded framework for neoantigen vaccine maturation

**DOI:** 10.1101/2024.08.14.607669

**Authors:** Guanqiao Zhang, Yaqi Fu, Kevin C. Chan, Ruofan Jin, Yuxuan Yang, Ruhong Zhou

## Abstract

Effective cancer immunotherapy hinges on the precise recognition of neoantigens, presented as binary complexes with major histocompatibility complex (MHC) molecules, by T cell receptors (TCR). The development of immunogenic peptide predictors and generators plays a central role in personalizing immunotherapies while reducing experimental costs. However, the current methods often fall short in leveraging structural data efficiently and providing comprehensive guidance for neoantigen selection. To address these limitations, we introduce NEOM, a novel neoantigen maturation framework encompassing five distinct modules: “policy”, “structure”, “evaluation”, “selection” and “filter”. This framework is designed to enhance precision, interpretability, customizability and cost-effectiveness in neoantigen screening. We evaluated NEOM using a set of random synthetic peptides, followed by available clinically-derived peptides. NEOM achieved higher performance on generated peptide quality compared to other baseline models. Using established predictors for filtering revealed a substantial number of peptides with immunogenic potential. Subsequently, a more rigorous binding affinity evaluation using free energy perturbation methods identified 6 out of 38 candidates showing superior binding characteristics. MHC tetramer peptide exchange assays and flow cytometry experiments further validate five of them. These results demonstrate that NEOM not only excels in identifying diverse peptides with enhanced binding stability and affinity for MHC molecules but also augments their immunogenic potential, showcasing its utility in advancing personalized immunotherapies.

## Introduction

Neoantigens are immunogenic peptides presented by the Major Histocompatibility Complex (MHC) – termed Human Leucocyte Antigen (HLA) in humans – are typically found on cells undergoing infection or neoplastic transformation^1,2^. These peptides are often derived from the proteolytic cleavage of mutated tumor-associated proteins, mediated by the proteasome and cytosolic peptidases. Post-cleavage, these peptides are transported into the endoplasmic reticulum (ER) by the Transporter Associated with Antigen Processing (TAP) complex. Within the ER, peptides undergo further trimming by resident aminopeptidases. The mature peptides then assemble with HLA I molecules, forming peptide-HLA (pHLA) complexes that are subsequently presented on the cell surface. This presentation is crucial for eliciting immune responses^3–5^. T cell receptors (TCRs) on cytotoxic CD8+ T cells recognize these complexes, facilitating critical immune functions, including cytotoxicity, B cell assistance, and cytokine production^6,7^. In the context of cancer immunotherapy, the development of personalized vaccines comprising these neoantigens or other immunogenic peptides is being investigated. Such vaccines are tailored to the unique tumor profile of an individual, aiming to amplify and diversify the repertoire of tumor-specific T cells^8–13^.

Neoantigen identification hinges on complex interaction involving peptides, MHC molecules, and TCRs, with the affinity of neoantigens for HLA molecules being a key step for cellular immunity. Mass spectrometry (MS) serves as an accurate method for profiling peptides presented on the cell surfaces; however, it inevitably samples across the entire cellular proteome^14^, posting challenge in specificity. Alternative methods, such as biochemical reconstitution assays for binding affinity, are constrained by the limited throughput of solid-phase peptide synthesis^15,16^. Despite the recent development of promising experimental tool, such as the EpiScan platform^17^, there remains a substantial need for effective computational predictors.

Traditionally, prediction models operated under the presumption that the individual amino acids contribute a fixed binding energy to a peptide’s overall binding affinity for MHC molecules, with these energies being additive to yield a cumulative binding affinity^18–20^. To investigate the mutual information among positions, a variety of computational techniques have been employed. These include methods based on decision trees, hidden Markov model and deep neural networks, with notable examples being NetMHC and NetMHCpan, which have significantly advanced the field^21–25^. In addition, to accommodate MHC variants beyond the human repertoire, NetMHCpan-2.0 has been developed^26^. Motivated by observation that an ensemble of predictions could outperform individual algorithms^27^, a combined predictor, Consensus, was developed within the Immune Epitope Database (IEDB)^28^. Progress in neural network applications has led to the refinement of NetMHC into NetMHC 4.0, especially optimized for an array of human MHC alleles, including HLA-A, -B, -C and -E^29^.

The utility of computational models for predicting pHLA binding is tempered by two primary issues concerning the original training data. Firstly, these datasets predominantly comprise binding affinity data, neglecting other biological parameters intrinsic to the pHLA binding process. This limitation has steered research efforts towards MS experiments for additional insights. In response, NetMHCpan-4.1 and NetMHCIIpan-4.0 were developed, incorporating MS-derived data into their training algorithms. Despite this integration, these updated versions exhibit predictive performances similar to their predecessors when benchmarked against class I epitopes^30^. MHCflurry 2.0 represents another advance, employing dual models to assess both binding affinity and antigen processing^31^. The second challenge involves the accurate representation of certain peptides, particularly when cysteine oxidation leads to the formation of disulfide-linked peptide multimers, which can confound experimental results for cysteine-containing peptides^32^. Recently, the EpiScan platform and the EpiScan Predictor (ESP) were developed to address this problem and significantly enhanced the prediction accuracy for peptides involving cysteine-modification^17^.

The sequence-based nature of the aforementioned predictive models is subject to constraints, primarily their inability to integrate structural details, a limitation largely due to the sparse availability of comprehensive structural database. Given the impracticality of experimentally solving a large number of atomic structures, researchers have resorted to computational techniques such as conformation sampling, empirical energy minimization^33^, and homology modeling, with applications like PANDORA^34^. In addition, the rise of deep learning methodologies in structure prediction is drawing significant attention^35–37^. However, the latest iteration of these computational advances, such as alphafold_finetune^38^, has not yet demonstrated a markedly superior performance in structure and binding prediction when compared to traditional models.

Moreover, the quest for highly immunogenic neoantigens demands the development of effective sampling methods to enhance the utilization of emerging predictive models^39–41^. AOMP program is designed by using attention scores generated by TransPHLA to optimize mutated peptides with higher affinity in the TransMut framework^40^. PepPPO^41^ represents another innovation, using reinforcement learning to identify target peptides for a particular MHC allele, with the efficacy of its selections being reinforced by the MHCflurry2.0 algorithm^31^. Nevertheless, relying solely on a single predictive model or on multiple models that draw from analogous datasets could potentially skew the peptide search process, leading to suboptimal outcomes.

To incorporate both sequence and structure data, as well as to apply effective sampling methods, we introduce a comprehensive neoantigen maturation framework, NEOM. This framework is designed to extrapolate individual peptide sequences to an expanded repertoire of variants. This expansion is achieved through a suite of algorithms, resulting in a curated antigen pool. NEOM is systematically structured into five modules: (1) a “policy” module, which employs algorithmic guidance to mutate peptide sequences; (2) a “structure” module, with PANDORA^34^ serving as its foundational component, to generate structural adaptations; (3) an “evaluation” module, drawing upon a repository of structure-based and knowledge-assisted criteria for assessing potential neoantigens; (4) a “selection” module, which makes informed decisions on which neoantigens to retain; and (5) a “filter” module, tasked with screening the output to ensure that only the most promising candidates are pursued. This integrative framework stands as a tool for personalized treatment, offering enhanced flexibility and efficiency. The conducted validation experiments highlight the candidate generation quality of NEOM in multiple aspects, including binding stability, binding affinity for MHC molecules, and immunogenicity. To understand NEOM’s architecture, we further conducted trials on the random synthetic peptides, followed by peptides derived from clinical data^42–47^ (Table S1), including glutamic acid decarboxylase 65 peptide segment 114-122 (GAD65_114-122_, VMNILLQYV)^42^. These trials revealed that whole maturation process was crafted to be customizable and interpretable, paving the way for informed rational designs in the future.

## Results

### NEOM: a novel framework for personalized neoantigen maturation

Our NEOM framework aims to systematically generate tailored antigen sequences for individual patients, considering their specific HLA types and unique peptide repertoire (Figure 1). The core of this process is an adaptive Monte Carlo Markov Chain (aMCMC) algorithm, which starts with an initial peptide sequence from an established pool, serving as the “reference”. This reference sequence undergoes structure generation in the “structure” module and evaluation via the “evaluation” module. Concurrently, mutations are introduced to this reference sequence, producing a set of mutant peptides (typically 10 peptides per iterative step). Each mutant forms its own pHLA structure and has a unique “loss score” (more below). The “selection” module then compares these loss scores. Peptides that satisfy the Metropolis criterion are pre-selected and added to the expanded peptide pool. The peptide with the lowest loss score becomes the new reference sequence, initiating the next recursive loop. This iterative process continues until no significant improvement in the loss score is observed, or the iteration criterion is met. Finally, the expanded peptide pool is filtered using the “filter” module to identify the most promising vaccine candidates.

**Figure 1.**
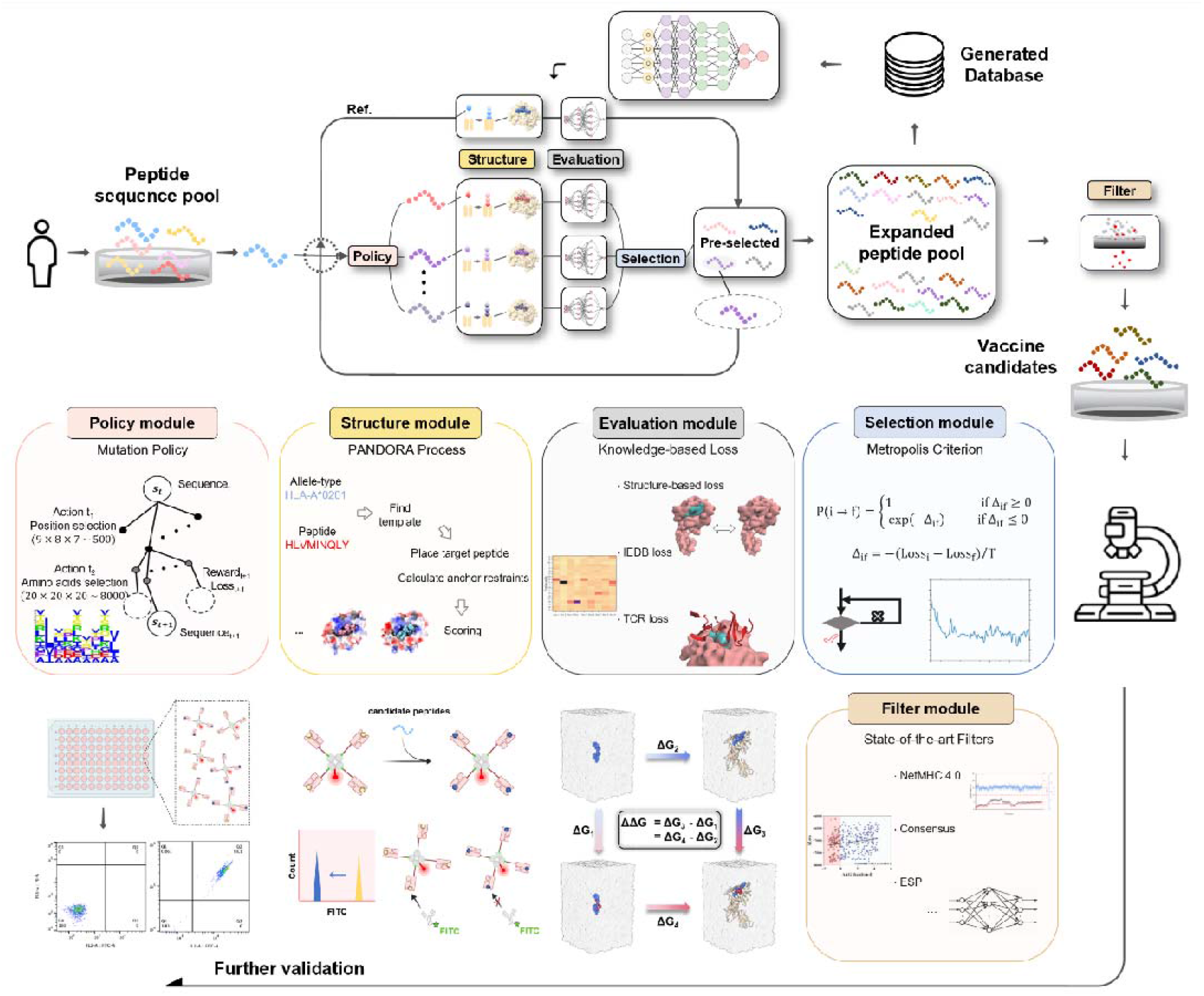
Neoantigen Maturation Framework NEOM. Commencing from a peptide sequence extracted from the patient’s original peptide pool, an adaptive MCMC optimization process entails successive iterations through policy, structure, evaluation, and “evaluation” modules. This iterative process culminates in an expanded peptide pool, followed by filtration through the “filter” module to obtain the most promising candidates. As for the module details, the figure elucidates the nuanced components within each module, which can be individually tailored. The “policy” module dictates the method by which peptide sequences are mutated. The “structure” module leverages structure generation tool to illustrate the pHLA structural intricacies. The “evaluation” module encompasses three segments: the Structure-based loss originating from the modeler, the IEDB loss, and the TCR loss. The “selection” module serves as the decision-maker, assessing the acceptance of mutants into the evolving peptide pool. All the sequences and corresponding loss scores could be collected as a generated database to train a neural network, replacing the “structure” and “evaluation” modules for acceleration. Additional computational prediction methods supplement NEOM as “filter” module, enhancing its efficacy in sieving results to yield highly reliable candidate peptides. These candidates can be further validated through various experiments.

### Assessment of the peptide generation quality

We use different predictors, such as NetMHC 4.0^29^, Consensus^28^, ESP^17^ and MHCflurry 2.0^31^ to characterize the quality of the expanded peptide pool of NEOM. The quality is assessed by better peptide rate, which defined as the binding affinity or immunogenicity of the peptide surpassing that of reference peptide within the expanded peptide pool. For instance, taking the peptide “VMN”, the glutamic acid decarboxylase 65 peptide segment 114-122 (GAD65114-122, VMNILLQYV)^42^ as a reference with HLA-A^*^02:01 as the model MHC type, we evaluate and compare with the baseline models. We compare three baseline methods: Random, IEDB+BLOSUM, TransPHLA-AOMP^40^ (details in Method). All these methods are independent of the aforementioned predictors. Among all results, NEOM achieved the rate of 0.518 in NetMHC 4.0^29^, 0.408 in Consensus^28^, 0.510 in ESP^17^, significantly outperformed other baseline methods (Figure 2a). Though in MHCflurry 2.0^31^, the rate of NEOM is slightly lower than that of TransPHLA-AOMP^40^, NEOM could generate more candidates than the latter. We also analyzed NEOM with different strategies, including different mutation policies based on prior knowledge, changing some hyperparameters and using neural network to replace “structure” and “evaluation” modules (details in Method), which obtained similar or even better results (Table S2).

**Figure 2.**
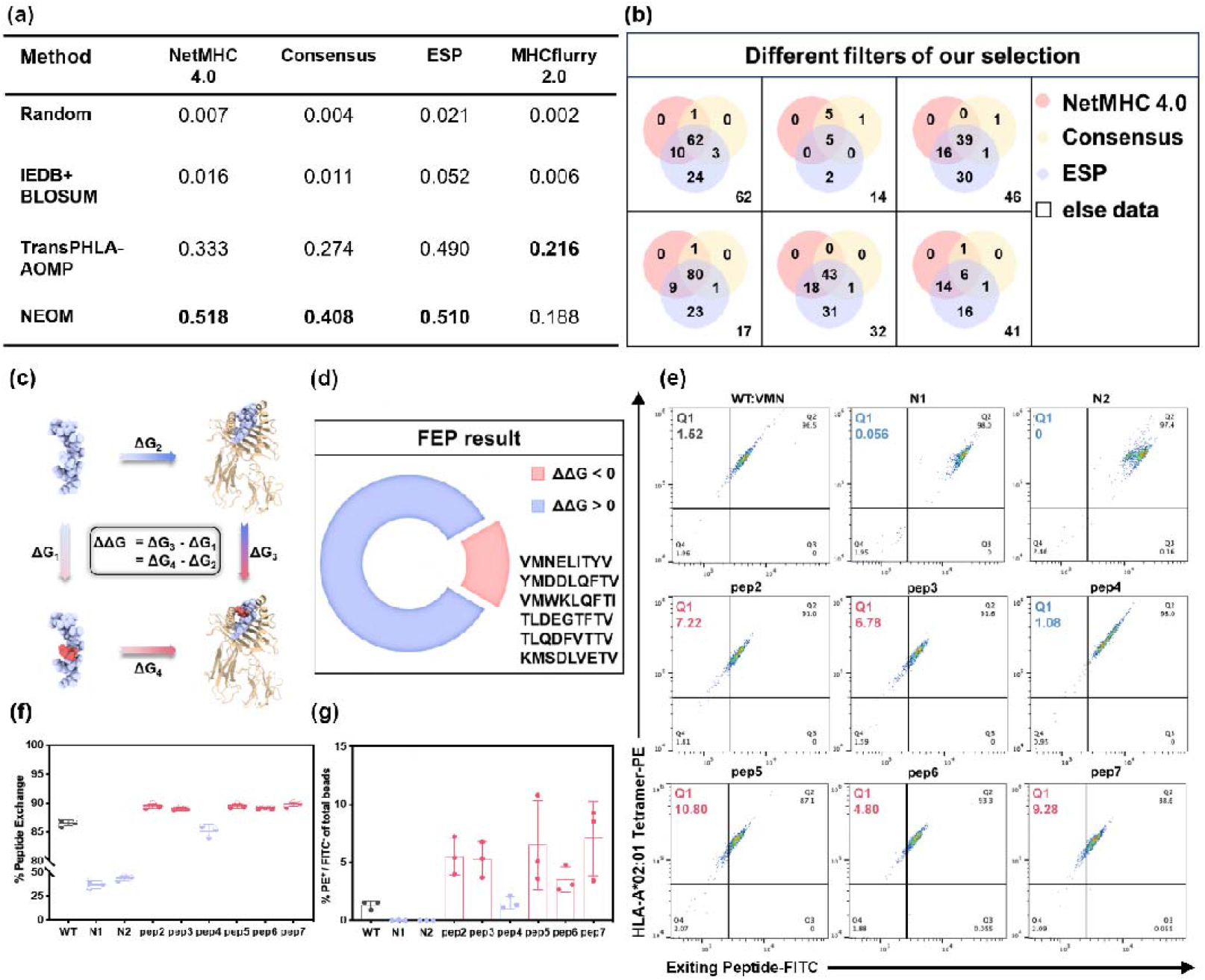
Performance summary and further validation with VMN to be reference peptide. **(a)** Assessment of the peptide generation quality using different state-of-art predictors, such as NetMHC 4.0^29^, Consensus^28^, ESP^17^ and MHCflurry 2.0^31^. **(b)** Outcome variations employing diverse filters across distinct optimization processes. **(c)** FEP procedure about the pHLA complex. **(d)** Results of the FEP about the candidate peptides, with 6 out of the total 38 selected peptides (roughly ∼1/6) exhibiting an enhanced binding to HLA-A^*^02:01. **(e)** Representative flow cytometry plots of HLA-A^*^02:01 tetramer^+^ and exiting peptide^-^. Numbers indicate percentages in quadrants. **(f)** The peptide exchange efficiency of wild type peptide and all mutant peptides. The experiment was repeated at least three times. Data are shown as mean ± SD. **(g)** Analysis of FCM showing the fraction of PE^+^/FITC^-^ of the total cells and the migration of the total population. It reflects the change in the mean fluorescence intensity of the population. The experiment was repeated at least three times. Data are shown as mean ± SD.

To make a thorough assessment of NEOM’s effectiveness, we employ a range of alternative filters in the “filter” module. Consider employing NetMHC 4.0^29^, Consensus^28^ and ESP^17^ separately to screen the expanded peptide pool. Optimal outcomes diverge based on the employed filter type (Figure 2b). However, the superior peptides occupied significant overlap, suggesting that the potential to identify valuable candidates for subsequent experimental phases remains robust even after extensive filtration.

### Computational and Experimental Validation

Machine learning predictors, despite their utility, are inherently limited by their dependency on finite datasets, particularly for biological systems. To address this issue, we utilized a combined validation approach with both computational and experimental techniques. We first examined the above predictions from deep learning with extensive all-atom molecular modeling. We performed standard molecular dynamics (MD) simulations to assess the structural stability of peptide-bound HLA complexes (Figure S1). Metrics such as peptide backbone RMSD, second residue (anchor residue) RMSD, and buried surface area were computed from the MD trajectories to evaluate the peptide’s inherent propensity to maintain HLA binding. Our results demonstrate that the majority of peptides can effectively bind to the HLA, with a success rate of ∼40% even with stringent criteria.

We further performed *in silico* mutagenesis studies with the free energy perturbation (FEP) method, which is regarded as the most rigorous and accurate method for free energy calculations^48,49^, to quantitatively evaluate the binding affinity of candidate peptides (Figure 2c). In FEP calculations, a reference peptide would be alchemically transformed into the target peptide (predicted candidates), during which the changes in binding free energies were calculated. Taking the VMN peptide as the reference, we have shown that 6 out of 38 tested peptides exhibit an enhanced binding to HLA-A^*^02:01 (Figure 2d). Subsequently, we performed peptide-HLA tetramer exchange assays (details in Method), in which the portion of a reference peptide being substituted by the candidate peptide were measured using flow cytometry. The analysis of FCM showed that five out of the six top candidate peptides bind substantially to HLA-A^*^02:01 (Figure 2e). The fraction of PE^+^/FITC^-^ population is proportional to the exchange efficiency of the candidate peptides. Gating strategy of flow cytometry was shown in the Figure S2. Therefore, we counted the proportion of PE^+^/FITC^-^ population to all beads (Figure 2g, Table S3). It can more clearly show comparison results between candidate peptides and wild-type VMN peptide. Notably, though the wild-type VMN peptide already showed a significant peptide exchange rate of approximately 86%, all our candidate peptides except mut4 (candidate #4) showed enhanced bindings (Figure 2f, Table S4). As negative controls, substituting anchor residues and additionally the charged residue at position 5 (N1, N2) with alanine significantly reduce the binding of tested peptides. Collectively, by integrating NEOM, molecular modeling, and experimental approaches, we successfully identified five peptides with optimal binding affinities and enhanced immunogenic potential. Our results also highlight the effectiveness of the NEOM framework in identifying peptides with enhanced binding stability and affinity for MHC molecules.

### Relationship between filter and evaluation modules

NEOM could provide a large amount of reliable vaccine candidates after integration of the “filter” module. This prompts us to explore the correlation between the “filter” module and other parts, especially our custom-designed “evaluation” module. Utilizing NetMHC 4.0^29^ as main predictor of the “filter” module, which is a widely used antigen scoring method, we acquired binding affinity results and compared them with the VMN peptide as reference to get ΔΔ*G*. Similarly, we calculated the differences of the loss function results in the “evaluation” module, comparing with VMN, to get Δ*Loss*. Red points are the peptides considered more valuable by NetMHC 4.0^29^ (Figure 3a). The resultant Pearson correlation coefficient of 0.22 suggests a mild correlation between Δ*Loss* and ΔΔ*G*.

**Figure 3.**
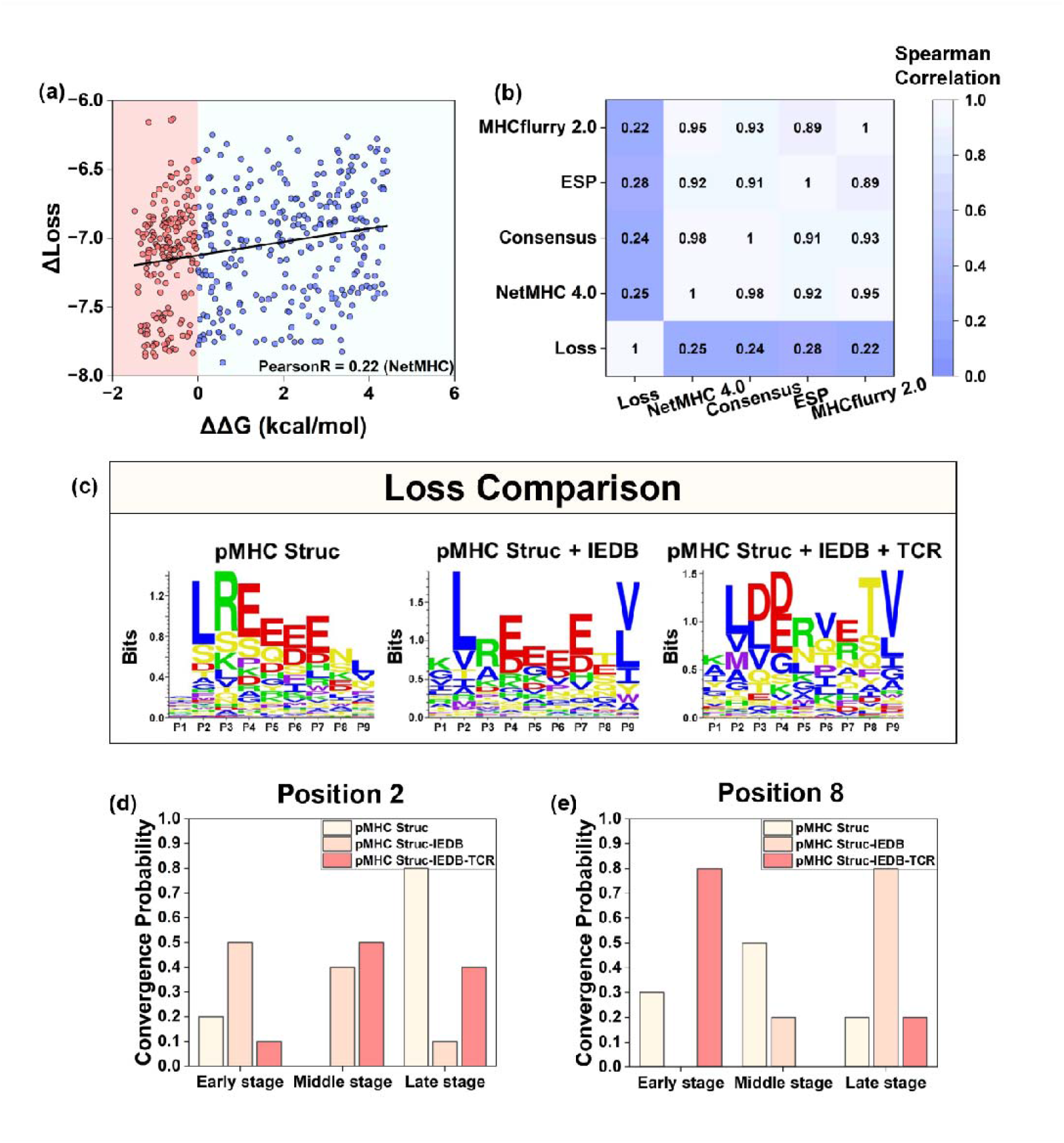
Deep interpretation of our designed loss function. **(a)** A slight correlation exists between Δ Loss and ΔΔG calculated by NetMHC 4.0. **(b)** Correlation heatmap between loss function and other predictors. **(c)** We conduct a comparative analysis of distinct residue probability distributions employing various loss functions. In this evaluation, the maturation process is undertaken ten times for each scenario, resulting in corresponding databanks. Each iteration involves 1000 optimization steps and employs a randomly selected start peptide. **(d, e)** Categorizing the initial 50 steps as the early stage, steps 50 to 250 as the middle stage, and the remaining steps as the late stage, we depict the probabilities of residues at positions 2 and 8 remaining unchanged within each respective stage.

Moreover, we use Spearman correlation to characterize monotonic relationship among the loss function and different predictors (Figure 3b). All predictors have high correlation coefficients of up to 0.9 between them, except for the correlation with 0.89 between ESP^17^ and MHCflurry 2.0^31^. Interestingly, the coefficients between the predictors and our designed loss function are no more than 0.3. This implies that our formulated loss offers an alternative approach to identifying immunogenic candidates, in addition to the traditional binding affinity metrics and machine learning based methods.

### The loss functions

To gain a deeper understanding of how this customizable loss function in the “evaluation” module influences the generated sequences, we analyzed residue probability distributions across multiple databases. For each database, we conducted 10 independent trials of the maturation process, with each starting from a distinct randomly chosen sequence and consisting of 1000 optimization steps. HLA-A^*^02:01 was chosen to be the model MHC type here. The resulting sequences exhibited intriguing patterns when varying the loss functions during the initial validation tests (Figure 3c). For instance, when solely relying on the pHLA structure-based loss function, we observed a significant dominance of charged residues in positions 3-7. However, incorporating the IEDB loss derived from Immune Epitope Database (IEDB) led to a noticeable increase in the probabilities of hydrophobic residues, particularly leucine (L) and valine (V), at positions 2 and 9. Those high frequencies of charged residues in positions 3-7, such as arginine (R) and glutamic acid (E), were slightly reduced. These changes underscore the substantial impact of the IEDB loss on the generated sequences.

To further refine the sequences for potential TCR activation, we integrated the TCR loss, derived from a TCR database informed by TCR-pHLA stability data from MD simulations (details in Method, Figure S3). This addition amplified the abundance of charged residues at positions 3-5, and 7, while nearly eliminating that at position 6. Interestingly, position 8 experienced a sharp rise in hydrophilic residues, particularly threonine (T). The probabilities of L and V at positions 2 and 9 remained dominant, but slightly attenuated. Given the fact that the alpha chain and beta chain in the complementary determining region (CDR) of TCR often interact with positions 3-8^50^, it is not surprising that the residue preferences in these positions were the most sensitive. Some charged residue preferences displayed significant increases when pHLA structure loss or TCR loss were biased, contrasting the pattern observed when only the IEDB loss was incorporated. These findings provide valuable insights into the basic interaction tendencies of TCR and highlight the importance of carefully selecting loss functions to guide the sequence maturation process.

### The Maturation Trajectories

To better understand how this loss function shapes the maturation landscape, we divided the process into three distinct phases: early (0-50 steps), middle (50-250 steps), and late (250-1000 steps) stages. This segmentation allows us to pinpoint when residues at specific positions stabilize and cease further alterations. Notably, we observed an accelerated convergence in the early phase (Figure 3d) when incorporating the IEDB loss alongside the pHLA structure-based loss. This acceleration aligns with our earlier findings, where we noted an increase in hydrophobic residue probabilities when IEDB loss was included. Similarly, introducing the TCR loss results in a faster convergence for position 8 (Figure 3e), also in consistent with our distribution analysis. Insights from other positions further support the notion that the maturation process is correlated to its outcome (Figure S4).

In parallel, we explore the effects of various mutation strategies on residue distributions and convergence propensities (Figure S5). While the results from these strategies are nonconclusive, to emulate the evolutionary process of mutation selection in nature, we selected the “frequency change with memory” method, maintaining a record of prior states.

### From synthetic peptides to real-world scenarios

To fully elucidate the mechanics behind peptide pool expansion through our methods, we analyze a randomly chosen peptide with HLA-A^*^02:01 again as the designated model MHC type. The maturation process, consisting of 1000 optimization steps in total, yields a swift decline in the loss function within just 50 steps, ultimately converging to a loss score of less than 1.5 (Figure 4a). Probability matrices for the 20 residues at distinct positions (Figure 4b-d) illustrate the evolutionary trajectory steering towards the local minima. Interestingly, the maturation routes could diverge, indicated by the acceptance events across different positions (Figure S6). For instance, both position 2 and 9 consistently start and end with hydrophobic residues, despite undergoing several mutations throughout the process (Figure 4e-f).

**Figure 4.**
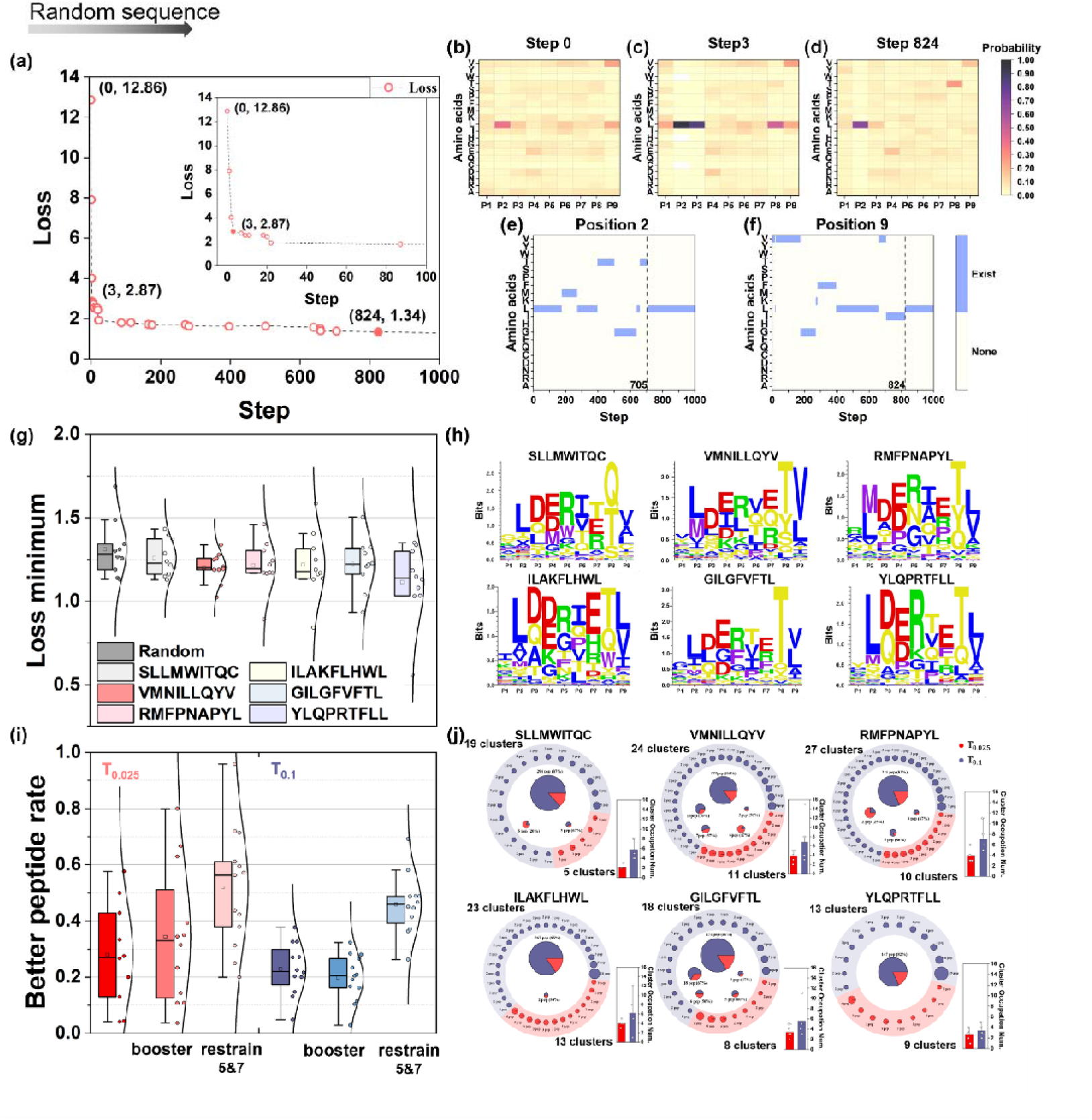
From synthetic peptides to real-world scenarios. **(a∼f)** Insights into the maturation process with a random start peptide. **(a)** A rapid decline in the loss score occurs within the first 50 steps, reaching a minimum loss value below 2000. **(b∼d)** Probability matrices for 20 amino acids in 9 positions. **(e, f)** Acceptance patterns for position 2 and 9. **(g)** Comparison of the loss minima from maturation trials using starting peptides sourced from real sequences versus peptides randomly generated. **(h)** Residue probability distributions from a database derived from 10 maturation trials, each initiated with a different peptide. **(i)** Evaluation of the improved peptide rate using NetMHC 4.0 as the criterion. Notably, results for T=0.1 demonstrate heightened stability compared to those for T=0.025, yielding a commendably comparable success rate with equivalent time expenditure. **(j)** Employing the clustering methodology offered by IEDB, we group the candidate peptides generated through five maturation processes with T=0.1 and T=0.025 using different starting sequences to discern the distinctions between them.

To interpret the method’s performance in more realistic setting, we applied it to different peptides from real-world sources (Table S1)^42–47^. Four of the peptides are derived from human and the other two are from virus, trying to ensure the multiplicity. Remarkably, the loss scores for these real peptides surpass most randomly chosen peptides (Figure S7). A quick comparison of the loss minima between randomly selected sequences and real peptides (Figure 4g) reveals that the averages of the loss minima for real peptides are slightly lower. To understand how the starting peptide may influence the generated sequences, we analyzed residue probability distributions of the corresponding expanded peptide pools. As for the VMN^42^, the methionine in position 2 has higher probability than in random starting sequences, similar to the “RMF” peptide (RMFPNAPYL)^44^. As for the “SLL” peptide (SLLMWITQC)^45^, the glutamine in position 8 has largest probability maybe due to its presence in the starting sequence. The “YLQ” peptide (YLQPRTFLL)^47^ shows a preference for threonine at position 6 and phenylalanine at position 7, while the “ILA” peptide (ILAKFLHWL)^43^ favors histidine at position 7. Interestingly, for the “GIL” peptide (GILGFVFTL)^46^, despite having valine at position 6 in the original sequence, the optimization process leads to a preference for threonine, highlighting unique characteristics in the starting peptide (Figure 4h). These results suggest that our method not only identifies real peptides as potent candidates but also effectively explores promising nearby minima of the starting sequences.

### Ways to discover more high-quality candidates

When the loss function is determined, ensuring a sufficient pool of candidates for subsequent validations holds great significance. To understand the influence when increasing determining factor of the pool’s volume, i.e. changing temperature T from 0.025 to 0.1 in the “selection” module, we assessed the better peptide rate through NetMHC 4.0^29^ as before (Figure 4i). For these two temperatures, we also provide some additional human-guided policies in the “policy” module, such as “booster” strategy and “restrain” strategy (details in Method), for efficient sampling. Evidently, the inclusion of these strategies considerably enhanced the quality of peptides. Despite a slight drop for the rates at T = 0.1, results demonstrate enhanced stability with more candidates compared to those at T = 0.025.

We next investigate the variation in breadth of expanded peptide pool when changing temperature. We utilized the IEDB clustering method^51^ to cluster the accepted peptides across five independent trials at two temperatures (Figure 4j). Clearly, a higher T value corresponds to a surge in cluster occupation number across all starting peptides, reflecting both augmented information capture and broader generalizability. Across sequences, the most dominant cluster consistently claims the most substantial fraction of all resulting candidates for both temperatures. Moreover, within the largest cluster, the contribution of peptides generated with T = 0.1 always surpasses that of T = 0.025. This implies that, within both temperatures, we can find this important local minimum near the starting point, yet T = 0.1, explores more. Regarding the smaller clusters, there appears to be no consistent pattern determining whether they will be explored simultaneously by both temperatures.

To determine the relative efficacy of these maturation results when changing the hypothetical temperature T from 0.025 to 0.1, we compared the cluster-to-peptide ratios of these two temperatures (Figure S8). Evidently, SLL tops the list, followed closely by VMN, indicating that the relative search efficiency (ranging from 0.9 to 1.0) remains largely stable even at elevated T values. While comparing the distinct clusters identified by the two temperatures for these sequences (Figure S9), SLL marginally lags behind VMN. For other peptides, even though their cluster-to-peptide ratios decrease, yet still remain above 0.5. These findings highlight the importance of identifying an optimal T range that balances the generation of a sufficient candidate pool with ensuring their validity. While higher temperatures can lead to more diverse candidates, excessively high temperatures can hinder convergence to desired minima (Figure S10). Overall, by carefully tuning the temperature parameter and using additional strategies, we can effectively balance exploration and exploitation in the NEOM framework, maximizing the chances of discovering high-quality peptide candidates.

## Discussion

In this study, we have developed a machine learning based framework NEOM, to enhance the efficacy of neoantigen maturation for future potential personalized immunotherapy. NEOM, characterized by its precision, interpretability, customizability and cost-effectiveness, leverages patient-specific HLA allele types and individual peptide sequences to generate a refined pool of antigenic peptides. The framework incorporates five key modules, namely “policy”, “structure”, “evaluation”, “selection”, and “filter”, each contributing uniquely to the neoantigen maturation process. As shown in assessment results, NEOM exhibited superior performance in generating peptides with enhanced immunogenic potential. Our “filter” module, utilizing predictors such as NetMHC 4.0^29^, Consensus^28^ and ESP^17^, has revealed that a considerable number of peptides with high immunogenic potential remain unidentified. With a more precise binding affinity evaluation based on rigorous free energy calculations, 6 out of 38 candidates are shown to exhibit favorable binding preferences to HLA (ΔΔ*G* < 0). Five of the top six candidates are further validated by peptide exchange and flow cytometry experiments. Notably, it demonstrates remarkable efficiency, capable of generating a suite of mutated peptides within approximately one minute on a typical GPU server. Our results demonstrate that the maturation process within this framework is fully visualizable, enabling detailed exploration of different modules and ensuring interpretability. The adaptability of the “evaluation”, “policy” and “selection” modules is a key feature, allowing for potential customization to meet diverse research needs. This finding underscores the efficacy of our approach in generating valuable antigenic sequences for future neoantigen based vaccine or other immunotherapy.

A critical aspect of NEOM is its adaptability, notably in the evaluation and “policy” modules, which have a variable impact on the mutational landscape. The impacts of these designs manifest not only the final outcomes, but also the rate of convergence. Various options exist for the “filter” module, especially machine learning predictors, showing high correlations with each other. We advise maintaining relative orthogonality between the customized loss function and the predictors in the “filter” module. One of the keys to activate the immune response is that pHLA can be presented by cells. Thus, our approach includes a structure-based loss component, enhancing the stability of the pHLA system and broadening the range of viable target candidates.

In our study, we acknowledge that solutions derived from a specific segment of the solution space may not be universally applicable. NEOM primarily targets peptides with high immunogenic potential. However, relying exclusively on predictors based on IEDB only can be limiting, as it represents only a fraction of the possible immunogenic peptides. To elevate the chance of identifying candidates, we approach this challenge from a different perspective – one that relates to, but does not overly depend on, the model based on IEDB. Central to our approach is the structure-based loss component, which significantly contributes to the stability of the pHLA system. This component is complemented by two statistical potentials derived from the IEDB and TCR databases, adjustable to individual immunological profile.

Finally, it should be noted that the versatility of NEOM extends beyond the HLA-A^*^02:01 allele, even though we use it as a model MHC for illustration here. When designing antigenic peptides, one common strategy is to expand the space of effective candidates before excessive and rigorous screening. Each module within NEOM is easily replaceable, setting it apart from traditional protocols. Our approach focuses on identifying an efficient maturation pathway to immunogenic antigens and explores potential candidates in their vicinity. Given the enormity of the vast sequence space, we recommend conducting multiple optimizations in parallel to navigate this space efficiently. While our current process is highly efficient, we have yet to incorporate reinforcement learning techniques to further this operation. Peptides and corresponding loss scores can be collected to train a neural network, but for different MHC types, we still need to generate enough data first. Another notable limitation is our reliance on the PANDORA-generated pHLA structures, which may not always correctly model the presented peptide conformations. As we look into the future, we aim to evolve NEOM into a proactive generator, incorporating additional structure-based and sequence-based strategies. We also foresee the integration of a deep reinforcement learning to enhance the speed and accuracy of NEOM. This advancement will involve a strategically designed sampling policy, aimed at converging to the minimum of loss function while also considering proximate candidate peptides.

## Methods

### NEOM architecture

The main goal of the NEOM architecture is to systematically identify antigenic peptide candidates for patients based on their unique neoantigens and specific HLA types (Figure 1). Once we determine the patient’s peptide sequence pool and corresponding HLA type, the optimization process commences. We initiate the maturation processes concurrently, using sequences from the original peptide pool as starting points. Central to this is the adaptive Monte Carlo Markov Chain (aMCMC), wherein one starting sequence serves as a reference point. To mirror real-world scenarios more closely, we either use a randomly generated peptide with a uniform distribution for residues in each position, or the sequences (Table S1) from the real world. Upon this selection, the corresponding pHLA structure of this reference is generated in the “structure” module and then evaluated in our “evaluation” module. Simultaneously, mutations are introduced to the reference sequence via the “policy” module, thereby producing a set of mutant peptides (typically 10 mutants each step). Following this, every mutant generates new pHLA structures, and the loss scores are calculated, with an average computation time of approximately 1 minute per step. Subsequently, a comparative analysis of these loss scores against the reference’s score takes place in the “selection” module. Mutants meeting the requests of the “selection” module are pre-selected to be part of the expanded peptide pool. Then, the mutant with the lowest loss score is regarded as new reference, executing a recursive loop. The iterative process is sustained until the standard deviation of the loss within the last 100 steps falls below a certain tolerance, or a predefined iteration limit (typically 1000 steps) achieved. After that, the “filter” module screens the resulting peptides to highlight the most promising ones as vaccine candidates. Further, these selected peptides undergo experimental validation to confirm their immunogenicity and assess their potential applicability.

### Policy module

To enhance the efficiency of the MCMC sampling process, the proposal distribution undergoes modifications to be an adaptive MCMC. Initially, mutation positions are chosen through a uniform probability distribution (Figure 1). Subsequently, the selection of mutated amino acids in corresponding positions occurs via a changing probability distribution, or probability matrix, and we mutate the amino acids of the specific positions for a given number (10 times each step). This distribution’s adaptability hinges on the current acceptance scenario and originates with a linear combination of equally weighted BLOSUM62 frequency and IEDB frequency that calculated from the Immune Epitope Database (IEDB). Basically, the sampling method about “Frequency change” means,

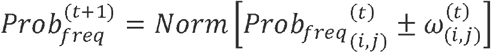

*t* means the optimization step, and 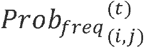 means that the probability of the residue *i* in the position *j* at *t* step. 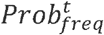 is a marginal probability that the sum of all the elements in one specific position is 1. There is also a parameter, 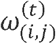, that adjusts the elements corresponding to the mutated residues of the probability matrix. *Norm* (·) is the normalization function. Additionally, here we can also use some other sampling methods. Usually, we would like to use “Frequency change with memory”, which means that the probability distribution will change with the probability of the history, just like the behaviors of the nature.

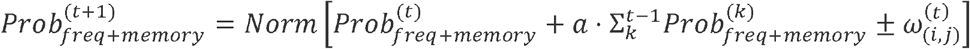

This 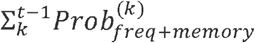 represents the history. And “No Frequency” means the probability will not change under the maturation process, only using the starting probability along the way.

Moreover, we also generate some specific restrains based on expert knowledges. In the “booster” strategy, we enforce that the mutant should have at least three same residues among position 3 to 8 (by default) when aligning with peptides in our TCR database. In the “restrain” policy, we impose a constraint to ensure some specific positions to be hydrophobic residues, such as maintaining positions 5 and 7 to be the hydrophobic residues.

### Structure module

To get more details about the pHLA system instead of only a sequence, PANDORA is employed for the pHLA structure generation. Inputting the peptide sequence and the specific MHC type, PANDORA undertakes the search for relevant structural templates, positioning the target peptides within the complex, and subsequently computes anchor restraints integral to the modeling procedure. When applying several mutated peptides in one step, it can generate all the structures simultaneously^34^.

### Evaluation module

The “evaluation” module comprises three distinct components (Figure 1). Initially, we adopt a modeller’s loss, combining both Discrete Optimized Protein Energy (DOPE) and molpdf score together, as a fundamental element of our knowledge-based loss to assess the structure of the pHLA^52^.

To enhance the integration of real-world insights, we formulate two additional loss functions: the IEDB loss and the TCR loss.

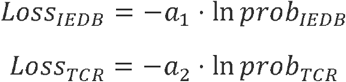

*prob*_*IEDB*_ is the probability that draws from sequences sourced from IEDB database, while *prob*_*TCR*_ is the probability that is calculated by the TCR-pHLA stability data, which consists of 52 peptides extracted from RCSB Protein Data Bank and determined to be stable through 100 ns all-atom molecular dynamics simulations.

These supplementary loss functions contribute a more comprehensive understanding of reality to NEOM.

### Selection module

The selection of the mutations obeys the rule, called Metropolis criterion, a well-established approach utilized for sampling from a specified distribution within a finite set (Figure 1). This criterion is a fundamental tool in computational physics and statistics, facilitating effective exploration of various possibilities.

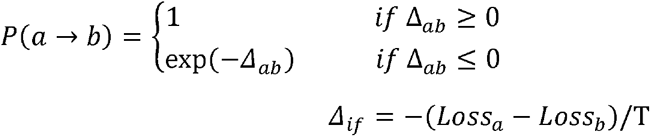

*P*(*a* → *b*)means the acceptance ratio from state *a* to state *b*, and T, the temperature, is a hyperparameter that could alter the number of accepted candidates. Remarkably, both settings for T = 0.1 or T = 0.025 consume roughly the same duration, predominantly governed by the processing time of the structure prediction step, with T = 0.1 displaying commendable success rates from a more expansive candidate list. Alternatively, one can also opt for the Metropolis-Hastings algorithm, offering a versatile method to sample from complex distributions.

### Filter module

Within this module, diverse filters can be used to scrutinize the expanded peptide pool, with the goal of extracting candidates with high reliability (Figure 1). Here we employ NetMHC 4.0 as a basic filter, with 0.5 and 2 to be the thresholds for strong binders and weak binders respectively, harnessing its sequence alignment capabilities based on artificial neural networks (ANN) to enhance the selection process^29^. Additionally, we present outcomes derived from other filters, including EpiScan Predictor (ESP), which adeptly avoids the underrepresentation of cysteine-containing peptides^17^, and Consensus, a method that combines predictions from various predictors to enhance overall predictive performance^28^. This comprehensive approach ensures a comprehensive evaluation of the peptide pool, refining the identification of promising candidates. The output comparison of these methods involved six unique maturation trials, each spanning 1000 steps (Figure 5c).

### Baseline models

To assess the generated candidate quality, we use three baseline models (Random, IEDB+BLOSUM and TransPHLA-AOMP^40^) for comparison. All these models are not developed based on the predictors for judging standard. Random is a method using uniform distribution probability to mutate the reference peptide. IEDB+BLOSUM is a method using uniform distribution probability to select a mutated position and use a combination of equally weighted BLOSUM62 frequency and IEDB frequency that calculated from the Immune Epitope Database (IEDB) to mutate the residue at that position. TransPHLA-AOMP is a method that optimize mutated peptides using attention scores generated by TransPHLA^40^.

### Sequence logo

To better understand the information content or significance of certain positions of the peptide sequences, a sequence logo is used to be the graphical representation. It consists of a stack of letters at each position, whose relative sizes of the letters indicate their frequency in the sequences. The total height of the letters depicts the information content of the position, in bits. The information content *I*_*j*_ of position *j* is given by:

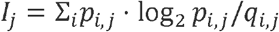

Where *p*_*i,j*_ and *q*_*i,j*_ are the observed probability (calculated from the database) and background probability, respectively, of the amino acid *i* in position *j*. Here we use Shannon sequence logo, then the height of the letter *i* in *j* column is:

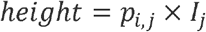

The lagger the letter *i* in *j* column, comparing with the other letters in the same column, the higher occurrence probability of *i*.

### Cluster method

To assess the exploratory capability when altering the important hyperparameters, we utilize the cluster tool that generates epitope clusters based on representative or consensus sequences^51^ in the Immune Epitope Database and Analysis Resource. By applying the “All the connected Peptides in a cluster” cluster method and 70% to be sequence identity threshold, we perform clustering on the sequences. This approach facilitates a clearer understanding of how changes in hyperparameters influence information extraction across peptide clusters.

### Correlation coefficients

The Pearson correlation coefficient, denoted as PearsonR, is a fundamental statistical measure utilized to quantify the linear relationship between two datasets. This coefficient offers insights into the extent to which changes in one variable correspond to changes in another. Specifically, it examines how the data points deviate from their respective means while taking into account the scales of the variables. This coefficient ranges between −1 and 1, where −1 signifies a perfect negative linear correlation, 1 indicates a perfect positive linear correlation, and 0 implies no linear correlation.

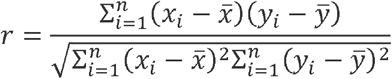

where *n* is the number of data points, *x*_*i*_ and *y*_*i*_ are the individual data points in the datasets x and y, respectively.

The Spearman correlation coefficient is a non-parametric measure of rank correlation. Unlike the Pearson correlation, which assumes a linear relationship, Spearman correlation assumes a monotonic relationship. It assesses the strength and direction of the association between two ranked variables. This coefficient also ranges between −1 and 1, similar to Pearson, but represents monotonic instead of linear correlation.

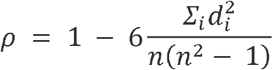

where *n* is the number of data points, *d*_*i*_ is the difference between the ranks of each pair of data points.

### Molecular Dynamics

We conducted all-atom molecular dynamics (MD) simulations on the complexes involving the peptides and HLA-A^*^02:01 systems. As for FEP, HLA-A^*^02:01-GAD65_114-122_ (VMNILLQYV) system was built from RCSB Protein Data Bank with PDB ID: 5FA3 and the intrinsic mimotope in the PDB structure was mutated to the GAD65_114-122_ antigen using VMD. The complexes were solvated in water boxes using the TIP3P water model. Na+ and Cl-were added to neutralize the systems and we set the ion concentration to be 0.15 M. Each system was energy minimized for 20000 steps, then equilibration period for 10 ns, followed by a 500 ns production run with a 2 fs time step (100ns production run for the structural stability test). All MD simulations were carried out with GROMACS using the CHARMM 36 force field. Long-range electrostatic interactions were calculated using the particle-mesh Ewald (PME) method and van der Waals (vdW) interactions were calculated using a cutoff distance of 1.2 nm. All production runs were performed with the NPT ensemble at 1 atm pressure and 310 K temperature. We further investigate trajectories sourced from MD simulations. This analysis primarily targets the structural stability of peptide-bound HLA complexes.

When investigating the structural stability of the complexes using MD (Figure S1), the structures are generated from PANDORA^34^. We undertake MD simulations of 100 ns across a spectrum of complexes. The peptide backbone RMSD (Pep-Backbone RMSD) emerges as a reliable metric to assess the peptide’s inherent propensity to maintain HLA binding. This metric alone yields an average success rate of around 80%, indicating that a majority of peptides can effectively bind to the HLA. Introducing a more rigorous definition of stability, one can incorporate the second residue RMSD (2nd-Res RMSD), buried surface area (Buried-SA), or a combination of both as additional criteria.

### Free Energy Perturbation

Following the MD simulations, the ultimate structure was chosen based on RMSD analysis, confirming structural stability (<0.5 nm) for over 200 ns. Multiple independent free energy perturbation (FEP) calculations were subsequently performed for each mutant. Residue mutagenesis investigations were carried out utilizing the FEP calculations. The binding free energy change Δ*G* caused by a mutation can be calculated as:

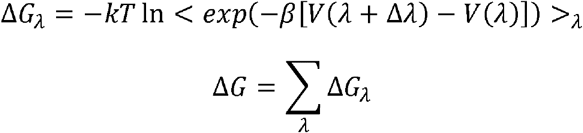

where *V*(*λ*) = (1 − *λ*) *V*_1_ + *λV*_2_, with *V*_1_ representing the potential energy of the wild type and *V*_2_ representing the potential energy of the mutant. When the system changes from wild type to mutant, the FEP parameter *λ* changes from 0 to 1, and < … > _*λ*_ represents average over the ensemble with potential *V*(*λ*). Given the long timescale of the binding process between two interacting surfaces, it is difficult to directly calculate the absolute binding affinity. To overcome this problem, we calculated the relative binding free energy change ΔΔ*G* with a thermodynamic cycle, as shown in (Figure 6a). This approach enabled us to derive the free-energy changes for the same mutation in both the bound and free states, thereby sidestepping the computation of intricate direct binding energies Δ*G*_*A*_ and Δ*G*_*B*_. In this thermodynamic cycle, the total change in free energy equals zero, and the relative binding affinity for the mutation from A to B is given by:

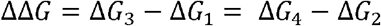

We conducted FEP calculations across 30 *λ* windows, with 600 ps per window, totaling 18 ns per calculation using a soft-core potential. Each FEP calculation was executed with no less than 3 replicas and then averaged to yield the final ΔΔ*G* value, culminating in a cumulative duration of at least 108 ns (18 ns x 3 replicas x 2 states, bound and free) for each mutation.

### Peptide-HLA tetramer exchange assays

All the peptides were synthesized according to standard procedure (Sangon Biotech, Shanghai, China). We used the QuickSwitch Quant HLA-A^*^02:01 tetramer kits^53^ (PE labeled) (Cat. TB-7300-K1, MBL International) to validate peptide exchange efficiency of all the potential antigenic peptides. The peptide exchange efficiency was used to compare binding affinity of in silico selected list of candidate peptides to HLA-A^*^02:01. Each peptide was dissolved in DMSO to a 1 mM solution to be assayed. Then, we pipetted 20 μL of HLA-A^*^02:01 tetramer into each well of the conical-bottom 96 well microtiter plate, added 0.4 μL of Peptide Exchange Factor plus 1 μL of peptide and mixed gently with pipetting. We repeated these steps for each additional peptide, including the Reference Peptide, and incubated the mixture 3 hours at room temperature protected from light. Next, the Magnetic Capture Beads were added to conjugated with the above tetramers, and the FITC-labeled Exiting Peptide Antibody was applied to the reaction against the Exiting Peptide. By measuring the percentage of original peptide replaced by a competing peptide, we may evaluate the exchange efficiency and compare the relative binding affinities of different peptides to HLA-A^*^02:01.

### Flow cytometry

In the Flow cytometric analysis, we detected the mean fluorescence intensity of FITC^54^. This assay reflects the relative binding affinity through MFI_FITC_, where the peptide exchange efficiency is inversely proportional to MFI_FITC_. We took the following procedure to quantify the peptide exchange efficiency: First, we ran control #1 sample (bead-captured QuickSwitch™ Tetramer) and adjusted compensation so that the MFI_FITC_ of bead control #1 equaled the MFI_FITC_ of the “Beads Only” control. Second, we ran control #2 sample, where beads had not captured any tetramer and therefore had no Exiting Peptide. The low MFI_FITC_ corresponded to 0% Exiting Peptide or 100% peptide exchange. Third, we ran control #3 sample, where beads had captured the QuickSwitch™ Tetramer. The high MFI_FITC_ corresponded to 100% Exiting Peptide or 0% peptide exchange. Then, we ran samples from well #4 and subsequent peptide exchange samples, noting the MFI_FITC_ of each. Peptide-exchanged tetramers will display various Exiting Peptide amounts, which are inversely proportional to the newly loaded peptide on the MHC molecules. Consequently, the measured MFI_FITC_ will be intermediary between MFI values obtained with bead control #2 and #3. Data analysis, gating strategy, and graphical presentation were done using FlowJo™ Software and shown in the Figure S2.

## ACKNOWLEDGMENTS

We thank Hong Zhou, Lei Fu, and Qinglu Zhong for helpful discussions. This work was partially supported by the National Key R&D Program of China (2021YFF1200404 and 2021YFA1201200), the National Natural Science Foundation of China (U1967217), the National Center of Technology Innovation for Biopharmaceuticals (NCTIB2022HS02010), Shanghai Artificial Intelligence Lab (P22KN00272), the National Independent Innovation Demonstration Zone Shanghai Zhangjiang Major Projects (ZJZX2020014), the Starry Night Science Fund of Zhejiang University Shanghai Institute for Advanced Study (SN-ZJU-SIAS-003).

## Author Contributions

R.Z. conceived the idea and designed the research. G.Z. wrote the software for five modules of NEOM with the support from R.J. G.Z. performed structure modeling and binding free energy calculations of the MHC–peptide complexes. Y.F. carried out the peptide-MHC tetramer exchange assay and flow cytometry experiments. G.Z., Y.F., K.C.C., Y.Y. and R.Z. analyzed the data and wrote the manuscript. All authors participated in discussions and revisions of the manuscript.

