## Supplemental Figures and Tables for "Multi-strategies embedded framework for neoantigen vaccine maturation"

Supplementary Information

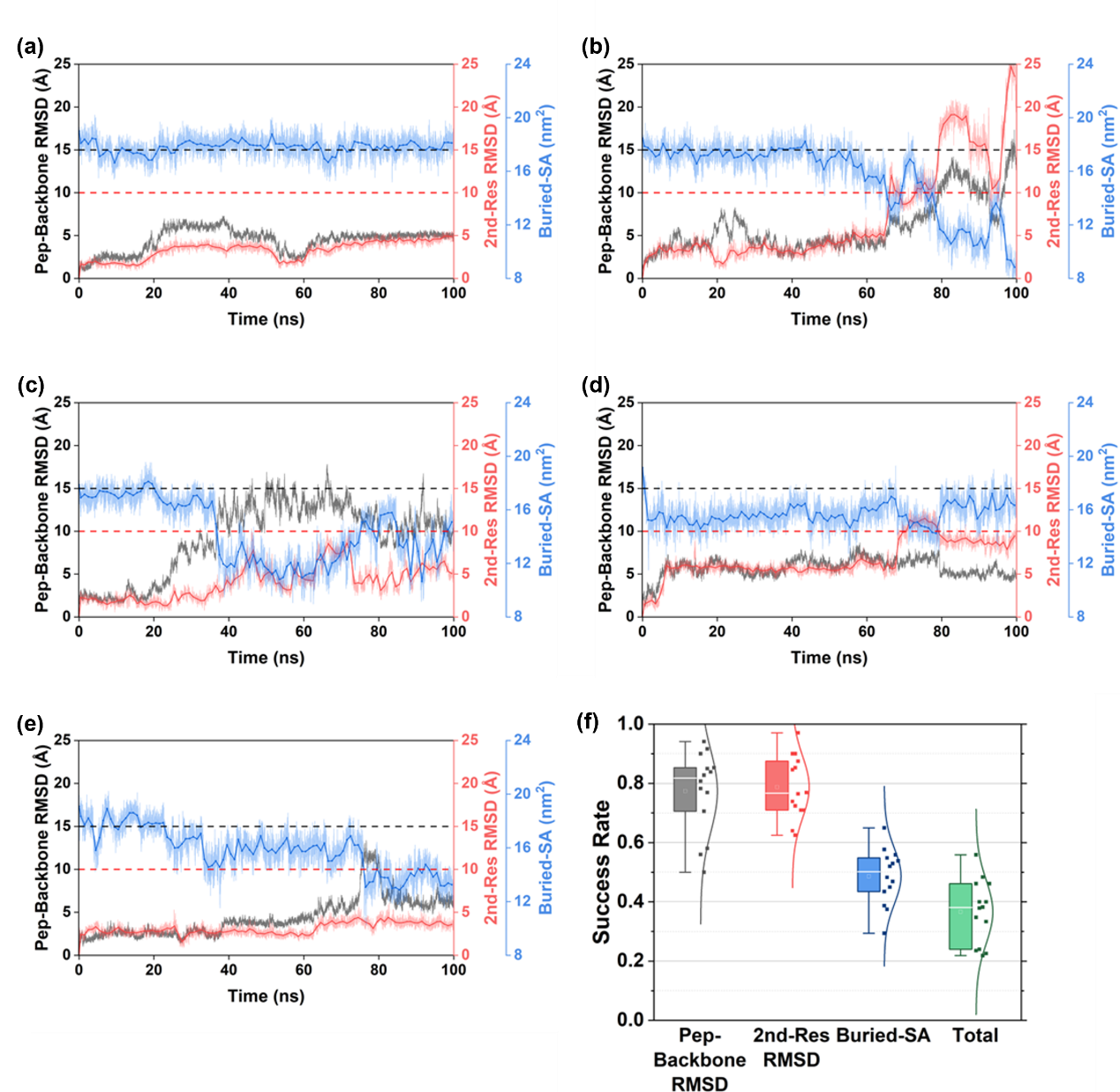

**Figure S1 Evaluation of the stable complex using various criteria**. Exploring the stability of the complex through a 100ns MD simulation, three key criteria are employed: peptide backbone RMSD (Pep-Backbone RMSD) < 15 Å, second residue RMSD (2nd-Res RMSD) < 10 Å, and the absence of a declining trend in buried surface area (Buried-SA). **(a)** Depiction of a trajectory conforming to all criteria. **(b)** Illustration of a trajectory violating all criteria. **(c~e)** Presentations of trajectories violating different criteria individually. **(f)** Success rates calculated utilizing distinct criteria. Remarkably, even with all these stringent criteria, the success rates consistently register at around 40%, underscoring the broad relevance and robustness of our optimization protocol.

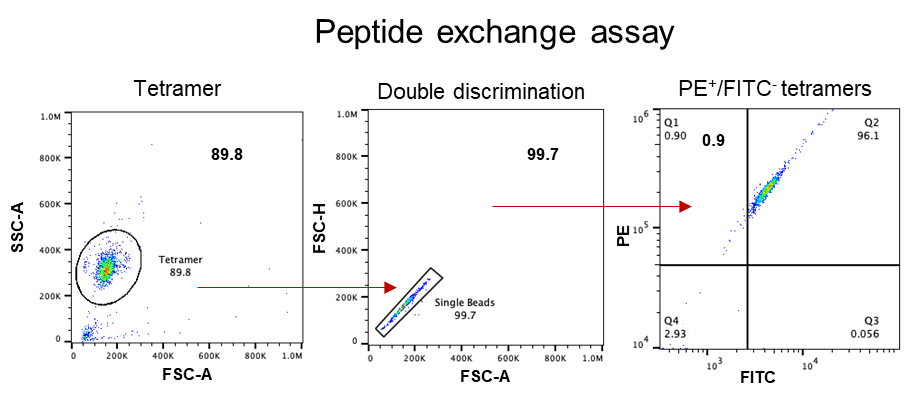

**Figure S2 Gating strategy of flow cytometry.**

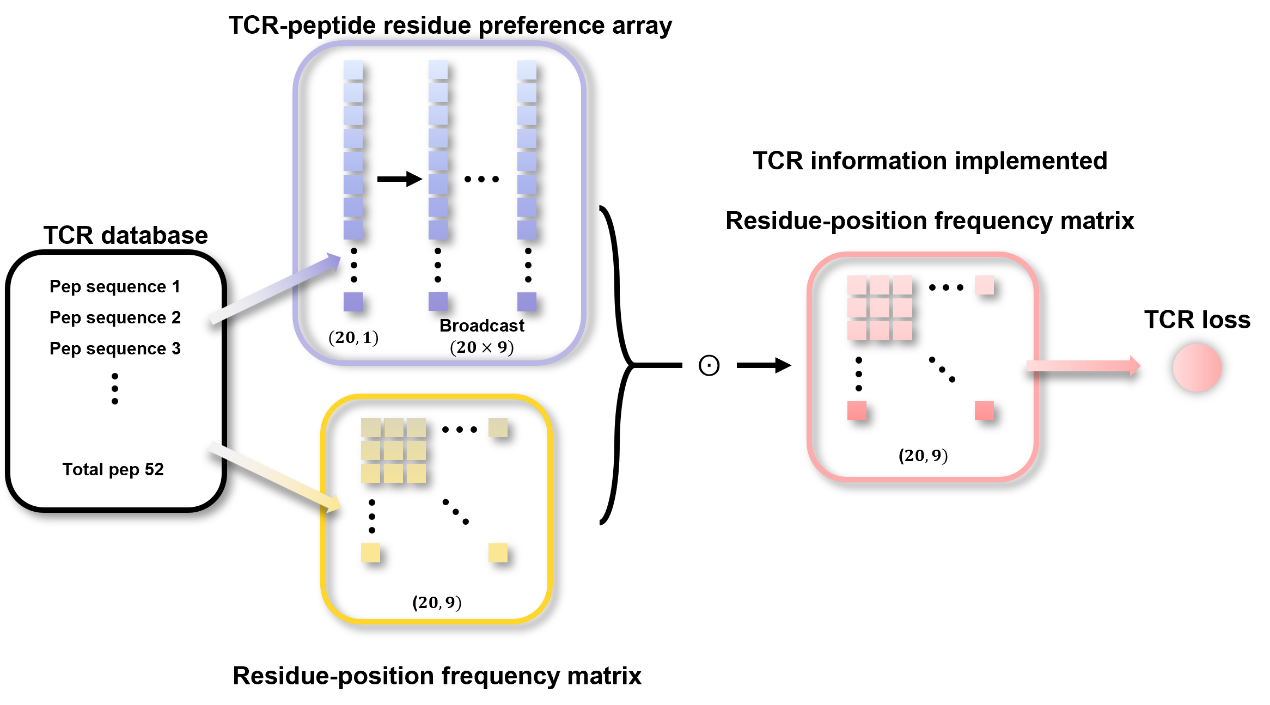

**Figure S3 Calculation of TCR loss.**

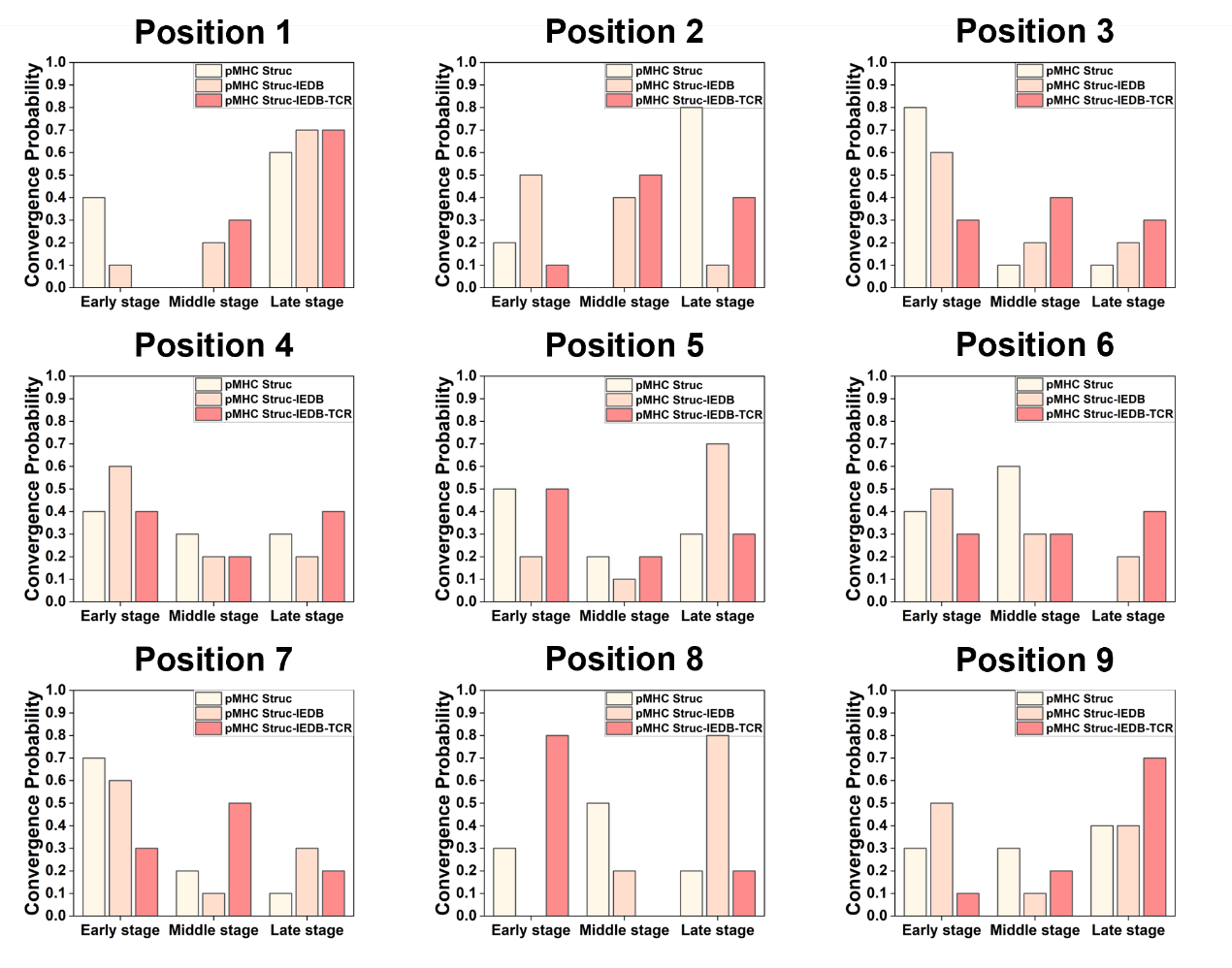

**Figure S4 Convergence probabilities with various loss functions.**

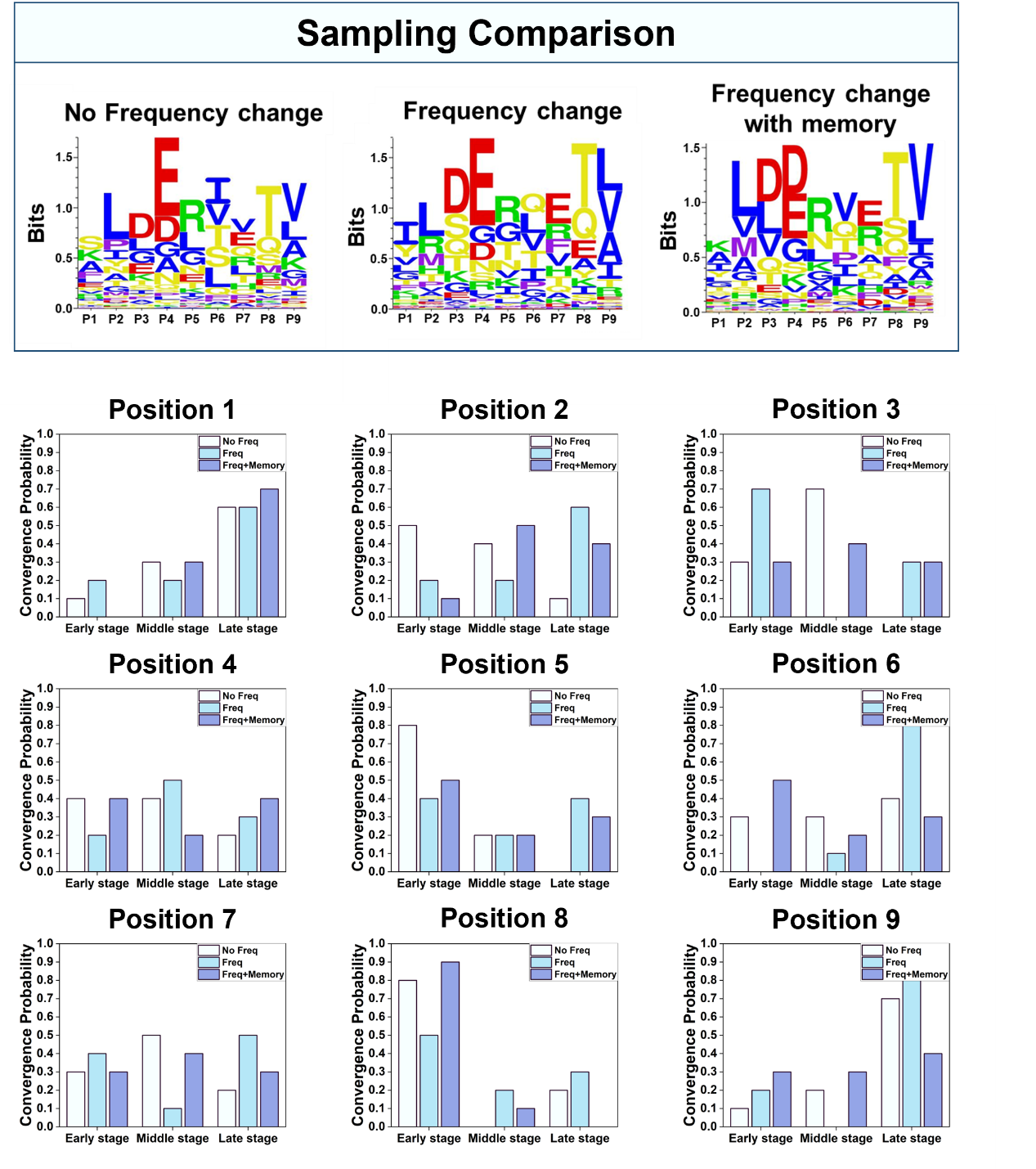

**Figure S5 Comparisons of outcomes with various searching methods.**

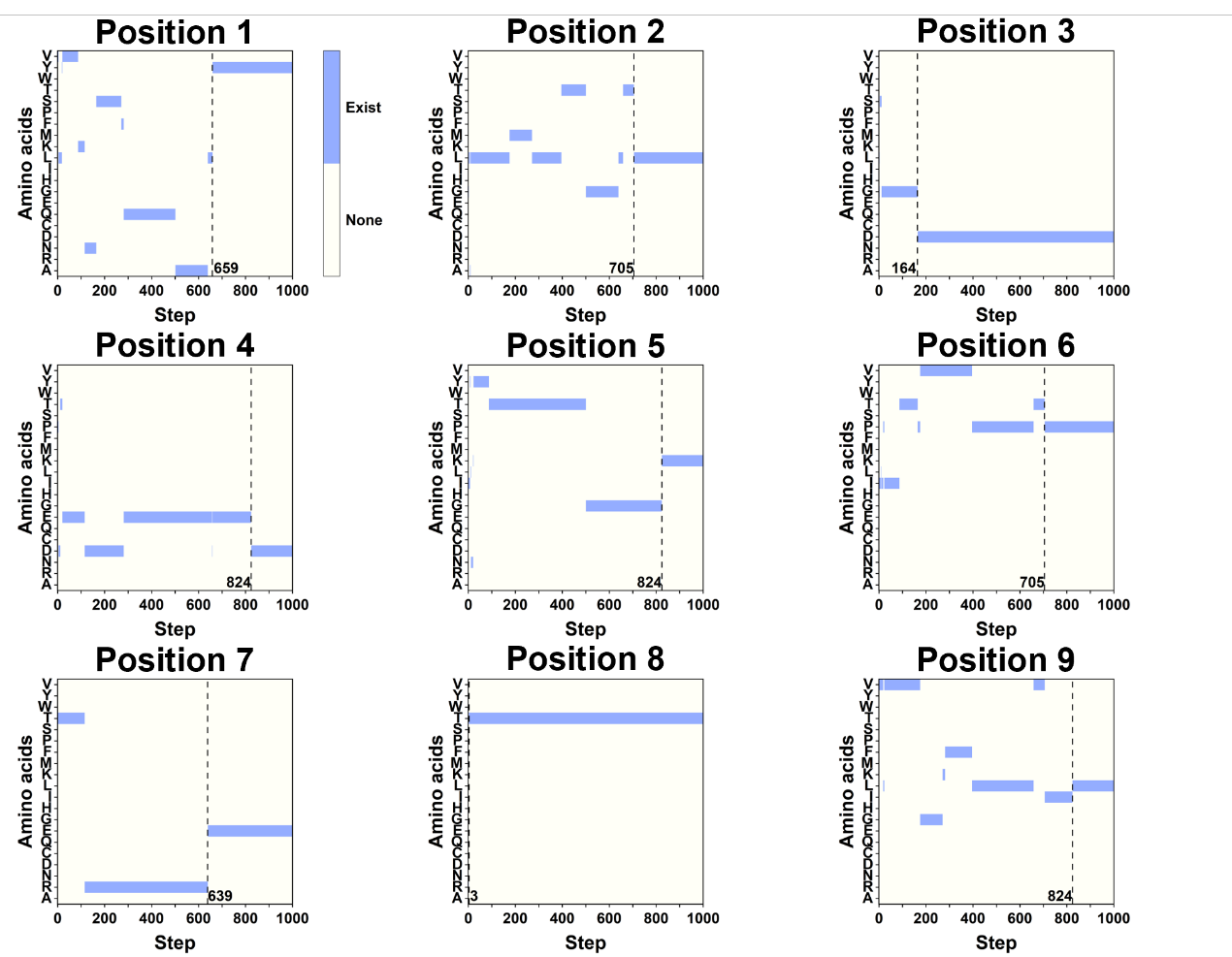

**Figure S6 Acceptance patterns for all positions with random start peptide.**

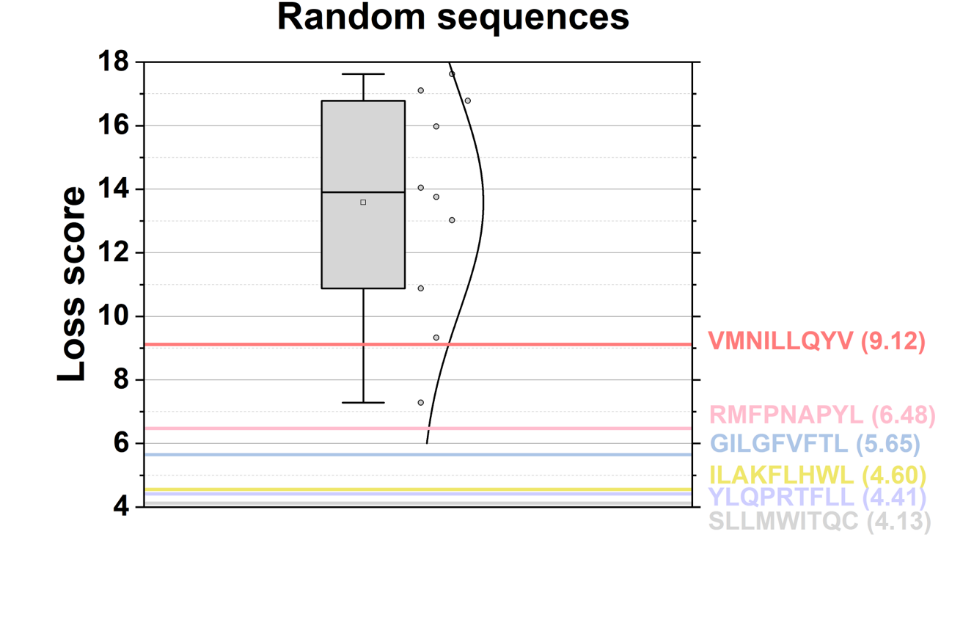

**Figure S7 Loss score for the random selected peptides.**

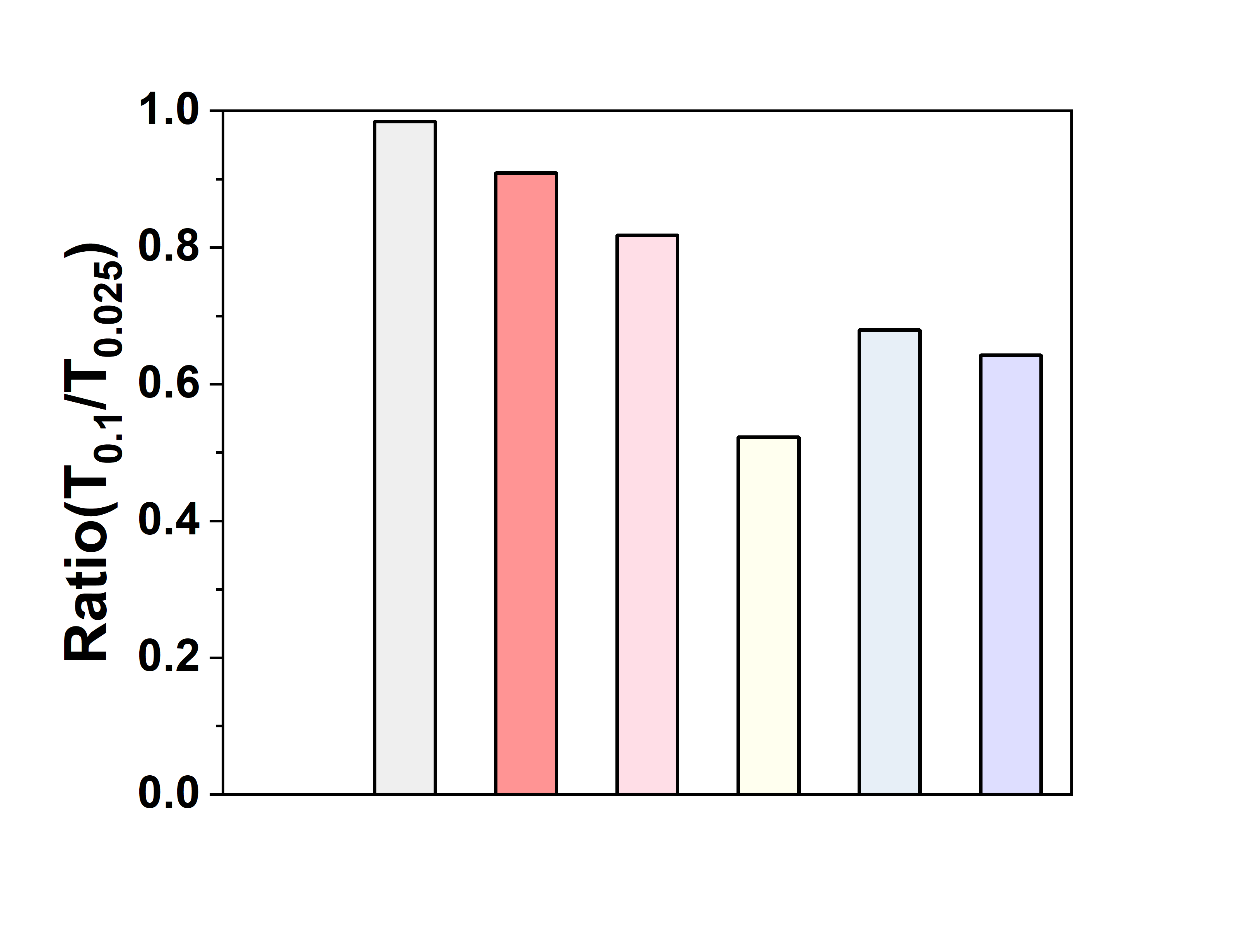

**Figure S8 Comparative ratios of "Cluster/pep" at** $\boldsymbol{T}\mathbf{=}\boldsymbol{0}\mathbf{.}\boldsymbol{1}$**versus** $\boldsymbol{T}\mathbf{=}\boldsymbol{0}\mathbf{.}\boldsymbol{025}$**for various initial peptides. "Cluster" denotes non-overlapping clusters, while "pep" refers to the total peptides we generated.**

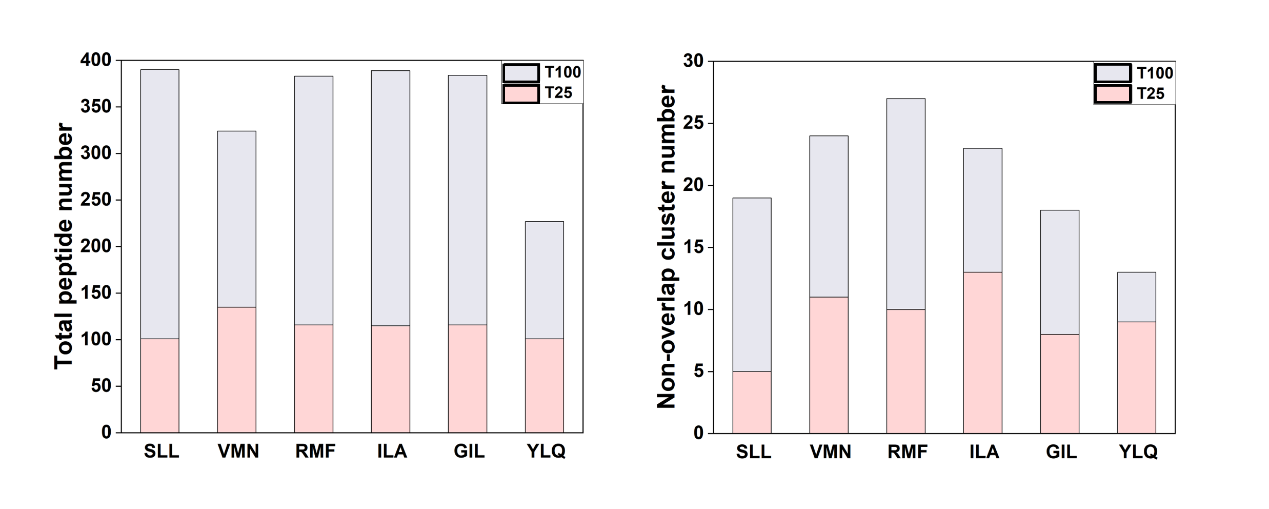

**Figure S9 Total peptide number and non-overlap cluster number of** $\mathbf{T=25}$ **and** $\mathbf{T=100}$**.**

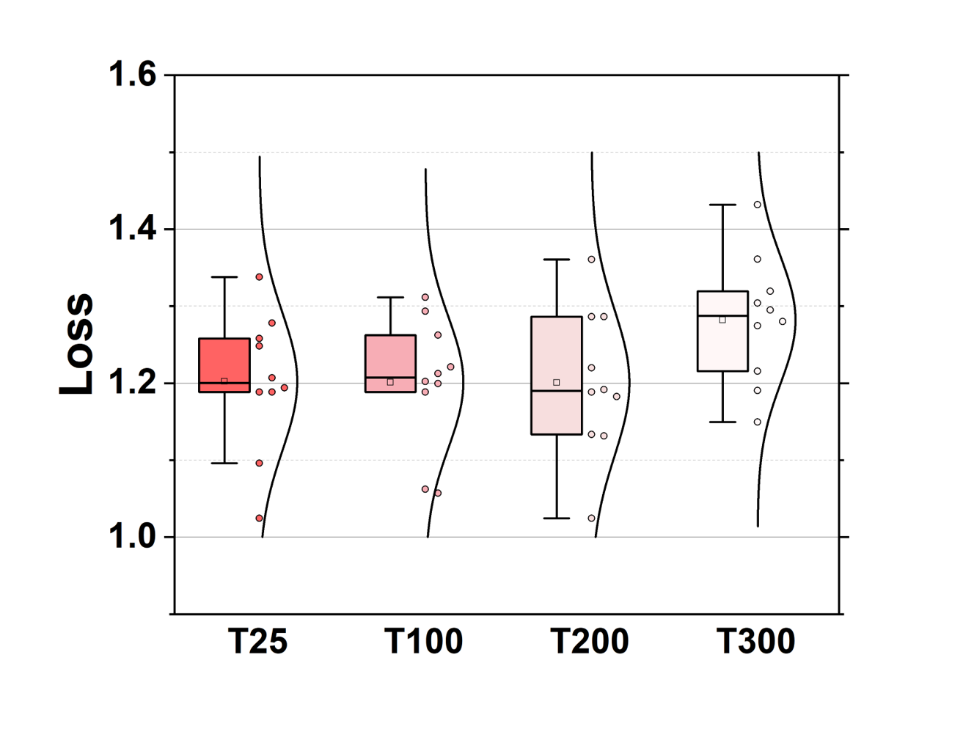

**Figure S10 Loss minimums with different temperature parameters.**

| **Sequence** | **Resource** | **Reference** |
| --- | --- | --- |
| VMNILLQYV | Glutamic acid decarboxylase GAD65_114-122_ | Knight et al., 2014 [40] |
| ILAKFLHWL | Human telomerase reverse transcriptase (residues 540-548) | Cole et al., 2017 [41] |
| RMFPNAPYL | Wilms tumor antigen WT_126–134_ | Holland et al., 2020 [42] |
| SLLMWITQC | pMHC tumor epitope NY-ESO_157–165_ | Sami et al., 2007 [43] |
| GILGFVFTL | influenza virus peptide MP_58–66_ | Stewart-Jones et al., 2003 [44] |
| YLQPRTFLL | SARS-CoV-2 spike protein epitopes (residues 269-277) | Wu et al., 2022 [45] |

**Table S1 Different sequences derived from the real.**

| **Method** | **NetMHC 4.0** | **Consensus** | **ESP** | **SMMPMBEC** | **MHCflurry 2.0** |
| --- | --- | --- | --- | --- | --- |
| Random | 0.007 | 0.004 | 0.021 | 0.004 | 0.002 |
| IEDB+BLOSUM | 0.016 | 0.011 | 0.052 | 0.010 | 0.006 |
| TransPHLA-AOMP | 0.333 | 0.274 | 0.490 | 0.255 | **0.216** |
| NEOM (T=0.1) | 0.229 | 0.190 | 0.448 | 0.183 | 0.065 |
| NEOM (T=0.1, booster) | 0.197 | 0.138 | 0.459 | 0.127 | 0.046 |
| NEOM (T=0.1, restrain 5&7) | 0.459 | 0.345 | 0.504 | 0.334 | 0.115 |
| NEOM (T=0.025) | 0.280 | 0.247 | 0.299 | 0.223 | 0.136 |
| NEOM (T=0.025, booster) | 0.344 | 0.252 | 0.522 | 0.250 | 0.170 |
| NEOM (T=0.025, restrain 5&7) | 0.518 | 0.408 | 0.636 | 0.412 | 0.188 |
| NEOM (T=0.1, net) | **0.557** | 0.438 | **0.807** | 0.417 | 0.091 |
| NEOM (T=0.025, net) | 0.553 | **0.487** | 0.713 | **0.479** | 0.175 |
| pepPPO (MHCflurry 2.0 based) | **0.709** | **0.628** | 0.728 | **0.627** | **0.485** |

**Table S2 Performance summary for the quality of generated peptides.**

is the model we show in the main text.

| **Peptide Sample/Sequence** | **% PE+/FITC- of total beads** | | |
| --- | --- | --- | --- |
| WT. VMNILLQYV | 0.900 | 1.520 | 1.520 |
| N1. VANILLRYA | 0.056 | 0.000 | 0.000 |
| N2. VANILLEYA | 0.000 | 0.000 | 0.000 |
| pep2.VMNELITYV | 4.010 | 7.220 | 5.410 |
| pep3.YMDDLQFTV | 6.780 | 3.690 | 5.360 |
| pep4.VMWKLQFTI | 1.340 | 2.130 | 1.080 |
| pep5.TLDEGTFTV | 3.600 | 5.110 | 10.800 |
| pep6.TLQDFVTTV | 2.680 | 3.100 | 4.800 |
| pep7.KMSDLVETV | 8.550 | 9.280 | 3.430 |

**Table S3 The PE+/FITC- proportion of wild type and other candidate peptides.**

| **Analyzed sample** | **MFI_FITC_** |
| --- | --- |
| Control #2: 0% Exiting Peptide or 100% peptide exchange | 321 |
| Control #3: 100% Exiting Peptide or 0% peptide exchange | 30567 |

| **Peptide Sample/Sequence** | **QuickSwitch MFI_FITC_ after Peptide Exchange** | | | **% Peptide Exchange** | | |
| --- | --- | --- | --- | --- | --- | --- |
| WT. VMNILLQYV | 4370 | 4224 | 4547 | 86.61 | 87.10 | 86.03 |
| N1. VANILLRYA | 19350 | 18143 | 20660 | 37.09 | 41.08 | 32.75 |
| N2. VANILLEYA | 18157 | 17039 | 16628 | 41.03 | 44.73 | 46.09 |
| pep2.VMNELITYV | 3522 | 3349 | 3578 | 89.42 | 89.99 | 89.23 |
| pep3.YMDDLQFTV | 3529 | 3698 | 3663 | 89.39 | 88.83 | 88.95 |
| pep4.VMWKLQFTI | 4710 | 4547 | 5153 | 85.49 | 86.03 | 84.02 |
| pep5.TLDEGTFTV | 3354 | 3518 | 3564 | 89.97 | 89.43 | 89.28 |
| pep6.TLQDFVTTV | 3622 | 3609 | 3550 | 89.09 | 89.13 | 89.32 |
| pep7.KMSDLVETV | 3539 | 3367 | 3304 | 89.36 | 89.93 | 90.14 |

**Table S4** The peptide exchange efficiency and MFI_FITC_ of all peptides.
